# Winner-take-all fails to account for pop out accuracy

**DOI:** 10.1101/2023.08.21.553875

**Authors:** Ori Hendler, Ronen Segev, Maoz Shamir

## Abstract

Visual search involves active scanning of the environment to locate objects of interest against a background of irrelevant distractors. One widely accepted theory posits that pop out visual search is computed by a winner-take-all (WTA) competition between contextually modulated cells that form a saliency map. However, previous studies have shown that the ability of WTA mechanisms to accumulate information from large populations of neurons is limited, thus raising the question of whether WTA can underlie pop out visual search. To address this question, we conducted a modeling study to investigate how accurately the WTA mechanism can detect the deviant stimulus in a pop out task. We analyzed two architectures of WTA networks: single-best-cell WTA, where the decision is made based on a single winning cell, and a generalized population-based WTA, where the decision is based on the winning population of similarly tuned cells. Our results show that WTA performance cannot account for the high accuracy found in behavioral experiments. On the one hand, inherent neuronal heterogeneity prevents the single-best-cell WTA from accumulating information even from large populations. On the other, the accuracy of the generalized population-based WTA algorithm is negatively affected by the widely reported noise correlations. These findings suggest the need for revisiting current understandings of the underlying mechanism of pop out visual search put forward to account for observed behavior.

## Introduction

The primary aim of visual search involves locating a specific object within a cluttered visual environment. Ensuring the organism’s survival demands both accuracy and speed in utilizing this ability, whether it’s for detecting food sources or pinpointing potential predators. In a visual search task, the object that the observer is searching for is termed the target, whereas the non-target items are termed distractors. Humans and other vertebrates perform different visual search tasks at differing degrees of efficiency which usually depend on the differences between the target and the distractors [1-3]. There is a consensus that there are two major search modes, known as parallel or pop out search and serial search [4-7]. These two search modes have been observed in humans, monkeys, archerfish, cats, and barn owls, thus illustrating the wide distribution of this visual behavior across vertebrate families [8-12].

The distinction between these two search modes is perhaps best illustrated in a classic experiment designed to assess search task efficiency where observers are asked to perform numerous search trials for an object while the number of distractors is varied. The time needed to produce the response, i.e., the reaction time, as well as the accuracy of the response are both measured.

In the pop out search mode, typical findings indicate that differences in visual features between the target and the distractors make the target more salient (Figure 1*A*− *B*) and lead to detection times that are independent of the number of distracting objects, as though the entire visual field were being processed in parallel (Figure 1*D*). Note that this rapid response time is associated with very high accuracy. In fact, success rates above 96% on a range of pop out tasks are common [13,14], with only a slight decrease as the number of objects is increased (Figure 1H) [15]. By contrast, in the serial search mode, the differences are less salient (Figure 1*C*), and no pop out is observed. In this case, the reaction time increases with the number of distractors (usually linearly, Figure 1*D*), and performance indicates serial visual scanning of the scene until the target is detected.

**Figure 1.**
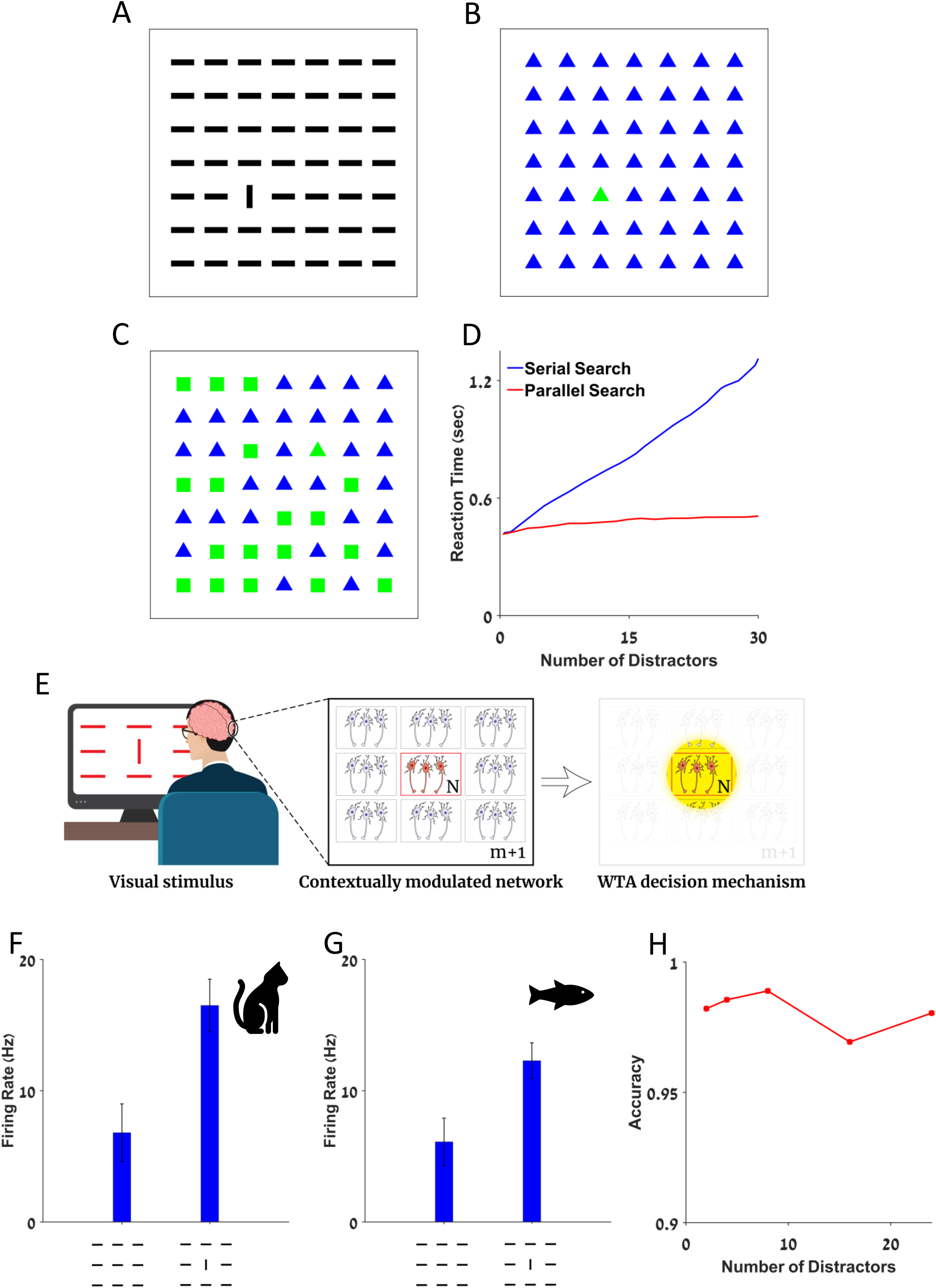
Pop-out, Behavior and Physiological Correlates. (**A-B**) Illustration of two pop-out stimuli: (**A**) A deviant vertical bar among numerous horizontal bars. (**B**) A deviant green triangle among many blue triangles. (**C**) A deviant green triangle among numerous green squares and blue triangles. (**D**) Reaction time of humans in serial and parallel visual search tasks, adapted with permission from [4] . (**E**) Schematic illustration of the model system. From left to right: Pop-out stimulus presented to the subject, network of *M* + 1 populations of *N* contextually modulated neurons each, WTA decision mechanism. (**F-G**) Example: mean firing rate of a single contextually modulated neuron in response to uniform and pop-out stimuli from the (**F**) visual cortex of a cat adapted with permission from [16] and the (**G**) optic tectum of a fish, adapted with permission from [17]. The dashed red line schematically depicts the classical receptive field of the neuron. (**H**) Success rate of Human subject in a pop-out task is plotted as a function of the number of distractors, adapted from [15].

The remarkable efficiency of pop out has prompted numerous experimental efforts to understand the underlying neural mechanism [18-21]. One widely accepted theory hypothesizes that pop out computation is implemented by a winner-take-all (WTA) competition between contextually modulated cells [22,23]. The basic architecture is illustrated in Figure 1*E*. These studies begin by the presentation of a pop out stimulus. In the figure the orientation of the bar is the stimulus variable (Figure 1*E*). In this case, the task consists of identifying the single deviant vertical bar from out of the set of horizontal bar distractors.

It is assumed that contextually modulated neurons respond to the pop out stimulus. These neurons are sensitive to objects placed outside their classic receptive field, i.e., they respond to the context of the stimulus. For example, data recorded in the cat and archerfish show that these cells fire at a higher rate when the stimulus within their classic receptive field is the deviant object than when the stimulus within the classic receptive field is a distractor (Figure 1*F* and 1*G*). Contextually modulated neurons have been found in the visual systems of primates, cats, birds, and fish [16,17,24-26].

In the next stage of processing, the WTA algorithm estimates the location of the pop out object from the location of the receptive field of the single most active neuron. It is generally assumed that since the visual scene is processed concurrently by the contextually modulated neurons, this phase is not sensitive to the number of objects. Next, the WTA reads the information from the entire population. The time required for this readout mechanism is inherently independent on the number of cells involved. Hence, the overall reaction time does not depend on the number of objects, as observed in behavioral experiments. On the theoretical level, the accuracy of WTA mechanisms have been studied in the framework of the two-alternative forced choice task [27]. The findings indicate that in a competition between two homogeneous populations, WTA accuracy increases slowly with the number of neurons in each population. The accuracy of a temporal variant of the WTA which estimates the stimulus based on the preferred stimulus of the fastest responding neuron exhibits dramatically deteriorating accuracy as the number of alternatives grows larger [28]. Thus, the weak ability of the WTA model to accumulate information from large populations of neurons and its difficulty addressing numerous alternatives raises doubts as to its ability to account parsimoniously for observed behavior in pop out tasks.

One possible solution is to consider a generalized WTA algorithm that estimates the target location by the population with the strongest response rather than by the single cell with the strongest response. However, it remains unclear whether this type of mechanism can yield the expected high accuracy reported in pop out tasks.

The goal of the study reported below was to examine whether WTA could serve as the mechanism underlying pop out in the visual system. To this end, we conducted a modeling study to investigate the performance of both the single-best-cell and a generalized-population WTA in a pop out task. We analyzed how various parameters such as the number of distractors, population size, contextual modulation strength, heterogeneity of the neuronal population, and noise correlations affected the ability of WTA to identify the deviant stimulus.

## Results

This section is organized as follows. First, we define a basic toy model for the stochastic neural response to a pop out stimulus and use this framework to analyze the accuracy of the single-best-cell WTA algorithm. Next, we apply the WTA algorithm to electrophysiological data of contextually modulated neurons, and analyze the effects of neuronal heterogeneity on the performance of the WTA. We then investigate the generalized WTA. We show that the generalized WTA emerges as superior in its ability to accumulate information from large populations but that this ability is undermined by noise correlations.

### Model architecture: single best cell setup

We consider a model of *M* + 1 columns, or populations, of *N* contextually modulated neurons each, responding to a pop out stimulus, Figure 1*E*. The pop out stimulus consists of one deviant object (in the receptive field of the ‘target’ column) and *M* distractors (in the receptive fields of the distractor columns, see example in Figure 1*E*). All neurons in a given column share the same receptive field and preferred feature (e.g., orientation, color, direction of motion, or shape). The neural responses, given the stimulus, are distributed independently. Neurons in the single target population that represents the deviant object fire with a mean response of *r*_*t*_ spikes. Neurons in the *M* distractor populations, of size *N*, fire a mean of *r*_*d*_ =*r*_*t*_/*q* spikes, where *q*>1 is the strength of the contextual modulation. The modulation ratio of contextually modulated neurons is an important parameter that serves as a proxy for the amount of information in the single neuron response. Empirically reported values of contextual modulation strength range from lower values of ∼1.1 to ∼1.5. We used this parameter in the analysis below. Specifically, we considered three types of distributions for the neural response: Poisson, exponential and Gaussian.

The task of the ‘readout’ mechanism is to identify the deviant (target) stimulus based on the neural responses. Next, we analyze the accuracy of the single-best-cell WTA, which we term WTA for brevity. The WTA algorithm estimates the target by the receptive field of the single neuron that fired the most spikes. In case of a tie, the WTA algorithm selects between the winners with equal probabilities.

### Winner-take-all performance in homogeneous populations

In the case of an exponential response distribution, analytical results for the dependence of the success rate, *P*_*c*_, on the number of distractors, *M*, and the modulation strength can be obtained in certain interesting limits. In the case of *N* = 1, that is, one neuron in each column, the accuracy of the WTA decision is given by (see Methods):

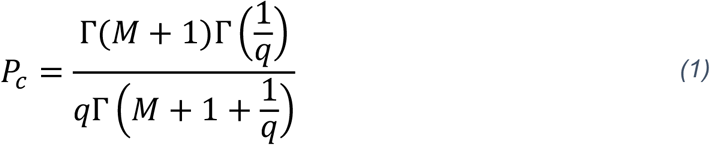

In the limit of *q* → 1, readout accuracy approaches chance value, 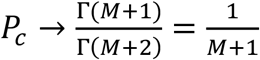, and for large *q* the accuracy converges to 1 algebraically in *q* (see Methods) as 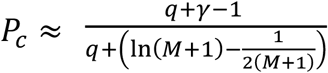. In the limit of a large number of populations 1 ≪ *M*, readout 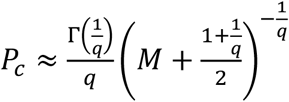accuracy converges to zero as *P*_*c*_ ≈ (see Methods). In the limit of large *N* and finite *M*, one obtains (see Methods):

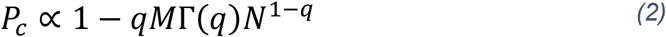

Thus, the probability of success of the WTA converges to 1 algebraically fast in *N* (Figure 2*A*). Nevertheless, the success rate decreases as the number of distractors, *M*, grows larger (Figure 2*B*). The only way to achieve independence with respect to the number of distractors is a ceiling effect. That is, when the performance approaches the maximal success rate of one, changes due to the number of distractors are small.

**Figure 2.**
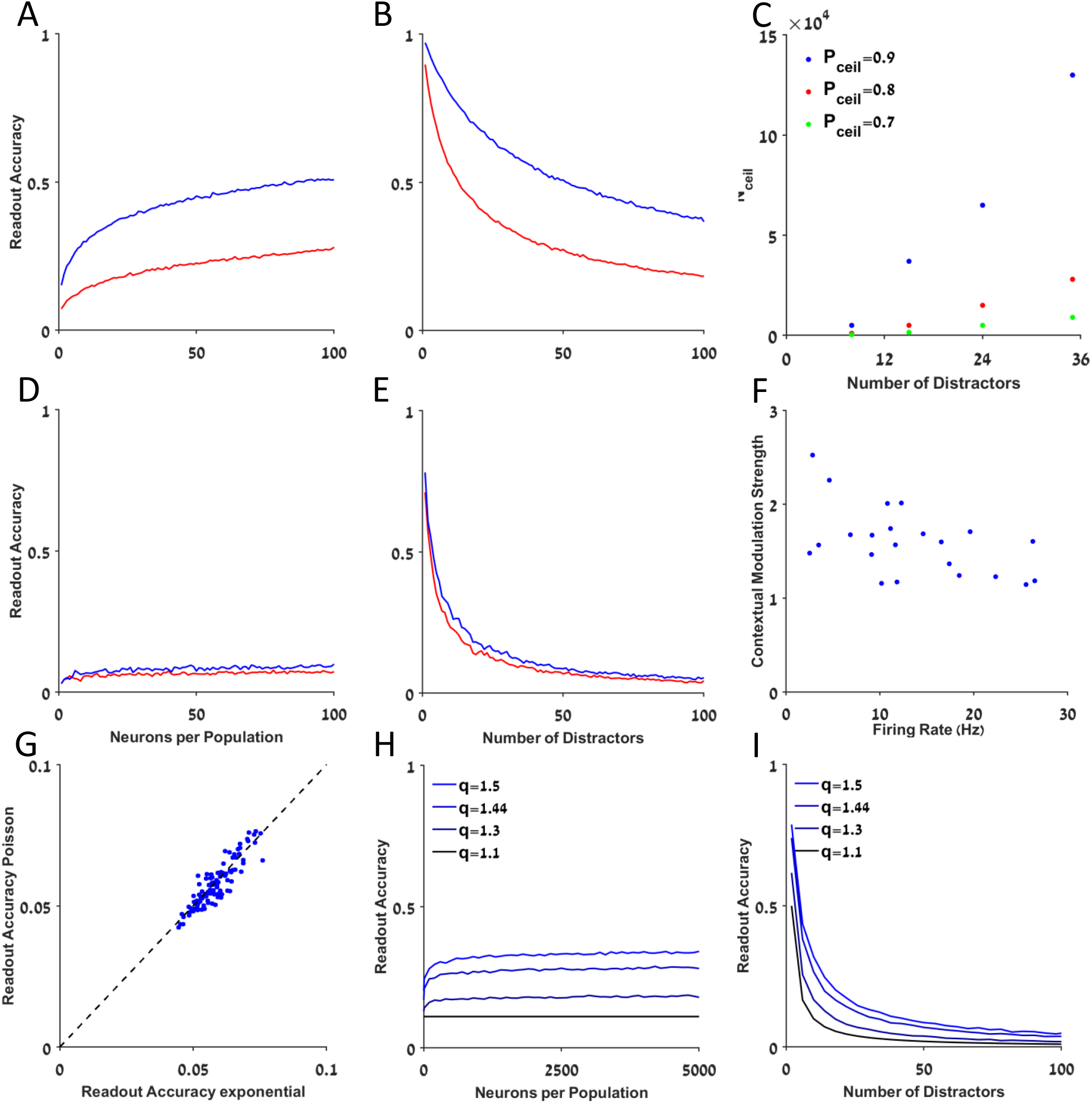
Accuracy of the single-best-cell WTA. (**A-B, D-E**) The accuracy of the single best cell WTA in homogeneous (**A-B**) and heterogeneous (**D-E**) networks is shown as a function of (**A, D**) the number of neurons, *N*, and (**B, E**) the number of distractors, *M*. The blue and red traces depict Poisson and exponential neuronal response distributions, respectively. (**C**) The number of neurons required to reach a certain accuracy threshold, *Nc*e*il*, is shown as function of the number of distractors, *M*, for different levels of accuracy threshold levels, *P*_*th*_, depicted by color. (**F**) Scatter plot depicting the response to pop-out stimulus and contextual modulation strength of 22 contextually modulated neurons in the optic tectum of the fish, the correlation between the two parameters is *ρ*=−0.53, *p* < 0.05, data adapted with permission from [17]. (**G**) WTA accuracy for Poisson population is shown as a function of its accuracy for exponential population, for the same realization of a neuronal heterogeneity, the correlation is *ρ*= 0.89, *p* < 0.05. The identity mapping is presented (dashed blue line) for comparison. (**H-I**) WTA accuracy in artificial heterogeneous networks is shown as a function of (**H**) the number of neurons, *N*, and (**I**) the number of distractors, *M*, for different contextual modulation strengths, depicted by color. The mean firing rates, *r*_*t*_, and mean contextual modulation strength, *q*, used in A-E, and G are the same and were taken from the data, F.

Specifically, denoting a tolerable level of error by *P*_ceil_, the ceiling effect can be achieved for *N* 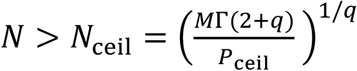 (Figure 2*C*). To account for empirical findings, one can select values of behavioral experiments in the range of *P*_ceil_ ∼ 0.95 − 1, *M ∼*10− 50 and values for *q* from electrophysiology *q* ∈ [1.1, 1.5]. Using these parameters, we found that *N*_ceil_ ≈ 10^4^ cells are needed to support the observed accuracy in the exponential case.

Qualitatively similar results were obtained for Poisson populations as shown in Figure 2*A* and 2*B*. In this case WTA accuracy decays to zero as the number of distractors, *M*, grows. In addition, its accuracy increases with the number of neurons per population, *N*, and a ceiling can be reached for *N* > *N*_ceil_ ≈ 10^4^.

### Single best cell WTA performance with a realistic architecture

Next, we analyzed the WTA algorithm with realistic parameters from neuronal recordings of contextually modulated neurons. The dataset consisted of *N*_data_ = 23 contextually modulated neurons from the optic tectum of archerfish responding to pop out and to uniform stimuli (Figure 2*F*) [17]. Each cell, *i* ∈ *{*1, … *N*_data_}, in the dataset was characterized by a pair of numbers: its mean response to a pop out stimulus, *r*_*i*_, (i.e., when the target is within its receptive field) and its contextual modulation strength, *q*_*i*_.

We first generated a realization of an individual animal (see Methods). To do so, we chose randomly in an independent manner, with equal probabilities and with repetitions *N* × (*M* + 1) neurons out of a pool of *N*_data_ neurons, 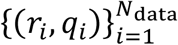. Each choice of *N* × (*M* + 1) neurons represent an individual animal.

Note that there are two types of randomness in our model. The first is the trial-to-trial fluctuations that result from the stochastic neural response. The other is the frozen or ‘quenched’ disorder that describes fluctuations between different individuals.

The accuracy of the WTA is depicted as a function of the number of neurons in each column, *N*, and the number of distractors, *M*, in Figure 2*D* and 2*E*, respectively. The accuracy of the WTA was estimated numerically by averaging over the trial-to-trial fluctuations for each given individual and then averaged over 100 realizations of different individuals.

Surprisingly, the accuracy of the WTA algorithm with a realistic architecture was considerably lower than in the homogenous case (compare Figures 2*A* and 2*B* with 2*D* and 2*E*), even though both shared the same average firing rate and contextual modulation strength. Furthermore, the rate at which realistic WTA accumulated information from large populations was also drastically reduced. The key difference lies in the fact that real neuronal populations are inherently heterogeneous, Figure 2*F*.

Interestingly, in the realistic WTA model, the effect of the neuronal response distribution (e.g., Poisson or exponential) on its accuracy was greatly diminished (red and blue traces in Figure 2*D* and 2*E*). Figure 2*G* depicts WTA accuracy for Poisson and exponential response distributions for the same realization of individuals. As can be seen from Figure 2*G*, even though the WTA accuracy was slightly higher under Poisson statistics, in general the accuracy under Poisson and exponential distributions was highly correlated (*ρ*= 0.89, *p* < 0.05).

### The source of the failure of single best cell WTA in the realistic model

To extend the analysis to a larger network we need to take into account the finite size of the empirical dataset. To overcome this issue, we generated artificial populations (see Methods) to study WTA performance for large heterogeneous networks which can mimic the essential features of the empirical data. We chose the parameters that characterized each individual from a model distribution with the same mean as the empirical data, and a variance equal to the square root of the mean. We found that the accuracy of the WTA increased monotonically in both the mean strength of contextual modulation and the number of neurons, *N*. Nevertheless, even for populations of *N* = 5000 neurons, the performance of the WTA was very poor (Figure 2H and 2I). For example, the readout accuracy was *P*_c_ ≈ 0.35 for *q* = 1.44 and *M* = 8 distractors, compared to the chance level of ≈ 0.11.

Thus overall, even for large networks, the accuracy was considerably lower than behavioral data suggests. In the example above, the WTA failed in more than 50% of the trials whereas the behavioral data were well above the 95% success rate.

To better understand the source of the poor performance of the WTA algorithm in realistic heterogeneous populations, below we briefly detour to examine what makes some individuals better than others. In a realistic model, WTA accuracy becomes a random variable that depends on the specific realization of neuronal heterogeneity. Figures 3*A* and 3*B* show the confusion matrix of the WTA algorithm for two individuals in a pop out task with *M* = 8 distractors for two extremes cases. The confusion matrix element, *C*_*i,j*_, is the probability of deciding the target location is at *i*, given that the target was at *j*. The diagonal, 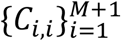depicts the probability of correct identification (the hit rate) for different target locations, *i* ∈ *{*1, … *M*}. For the first individual, Fig 3A, the hit rate varied from 0.2 to 0.6 depending on the target location. In the second example, Fig 3B, the hit rate was in the range 0.2 to 0.35. Thus, there was variability in the performance across locations and across individuals. Furthermore, we found that the hit rate of target *i, C*_*ii*_, was correlated (*ρ*= 0.76, *p* < 0.05) with the probability of erroneously estimating *i* to be the target, 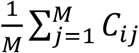, see Figure 3*C*.

**Figure 3.**
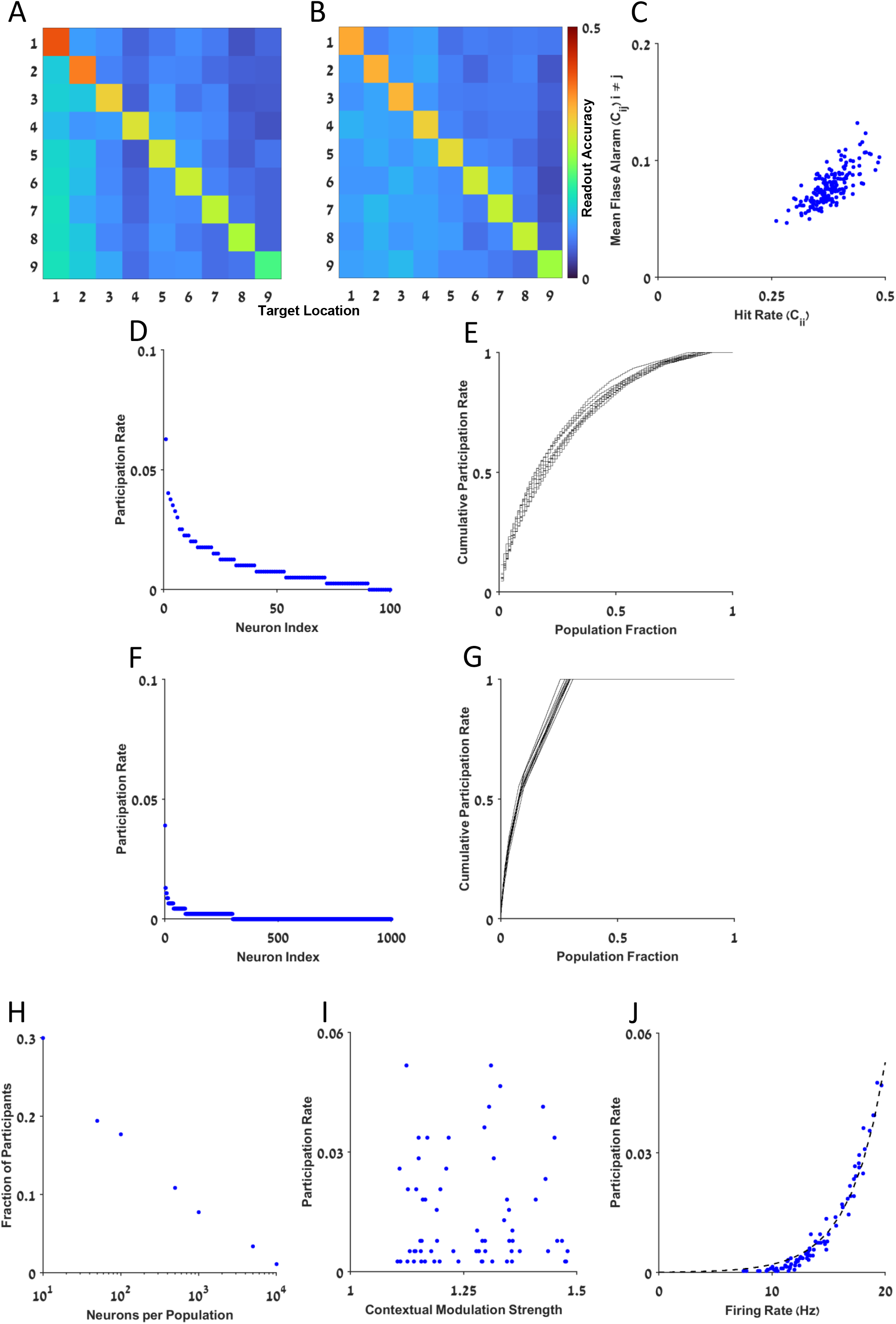
Investigation of WTA errors. (**A-B**) Two examples of confusion matrices presenting the performance of the WTA algorithm in in two example realistic (heterogeneous) networks. (**C**) The false alarm is shown as a function of the hit rate for different realizations of the network heterogeneity, correlation *ρ*= 0.76, *p* < 0.05. (**D**) Participation rate of different neurons is shown for one example of a single individual with *N* = 100 neurons per population (**E**) The cumulative sum of the participation rate of the *n* neurons in (D) with the highest participation rate, as a function of the fraction of neurons from the entire population, *n*/*N*. (**F**) As in D with *N* = 1000 neurons per population (**G**) As in E, for the neurons in F. (**H**) The fraction of neurons required to reach cumulative participation rate of 50%, is shown as a function of the number of neurons per population, *N*. (**I**) Participation rate of different neurons is shown as a function of their contextual modulation strength, *ρ*= 0.05, *p* = 0.67.(**J**) Participation rate of different neurons is shown as a function of their firing rate. The solid black line depicts exponential fit of the form *f*(*x*) = *ax*^*b*^, with, *a* = 1.77 · 10^−9^, *b* = 5.74 and *R*^2^ = 0.98.

What makes certain populations better than others at identifying targets? To shed light on this question, we examined which single neuron was responsible for the decision. In the WTA algorithm, a correct identification is said to occur when a single neuron in the target population is the winner. A high hit rate is expected when the decision is dominated by the activity of the more informative neurons, i.e., neurons that are characterized by high *q* values.

We defined the ‘participation rate’ of a neuron as the probability that the neuron was the ‘winner’, given a correct decision (see Methods). In the simplest case of a homogeneous population, every neuron has the same probability of being the winner, and the participation rate is expected to be 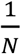.

In realistic non-homogenous cases, the participation rate is highly non-uniform. For example, Figure 3*D* depicts the distribution of participation rates in a heterogeneous population of *N* = 100. Here, seventeen neurons (17% of the population) were responsible for ∼50% of the decisions (Figure 3*E*). In another example (Figure 3*F*) with *N* = 1000 neurons, fewer than 70 neurons that made up 7% of the neural population were responsible for 50% of the decisions Figure 3*G*. Thus, a small fraction of the population was responsible for most of the decisions, and this fraction decreased as *N* grew larger, Figure 3H. For example, for *N* = 10000 neurons, barely 1% of the population made 50% of the decisions.

What characterizes neurons with a high participation rate? The analysis showed that the participation rate was not correlated with the contextual modulation strength (*q*) which is a proxy for the information content of single neurons (Figure 3*I*, *ρ*= 0.05, *p* = 0.67). Instead, neurons with the highest participation rate were the neurons with the highest mean firing rates, Figure 3*J* (*ρ*= 0.89, *p* < 0.05).

However, in empirical data, these two characteristics of the neuronal response tend to be either uncorrelated or negatively correlated, as was shown for the archerfish (Figure 2*F*, (*ρ*=−0.53, *p* < 0.05). Hence, the source of the poor performance of the WTA in realistic heterogeneous networks is that the WTA algorithm estimates the target in terms of the most active neurons, which are (roughly) uncorrelated with the most informative ones.

### The generalized winner-take-all architecture

One natural way to remedy the single best cell WTA algorithm is to consider competition between populations rather than between single neurons. In this generalized WTA competition, the stimulus is estimated by the most active population; i.e., by the population with the highest mean spike count across neurons. Specifically, the algorithm first averages the spike counts over all the cells in the population (or columns in Figure 1*E*), and this result is fed into the competition mechanism that selects the winner population.

Figure 4*A* presents the result of the readout accuracy in the generalized WTA algorithm for heterogenous populations. It shows that the generalized WTA accumulated information from large populations at a much faster rate than the WTA (compare Figure 4*A* to Figure 2*D*). This algorithm can achieve an accuracy of 95% using only a few hundred neurons per population.

**Figure 4.**
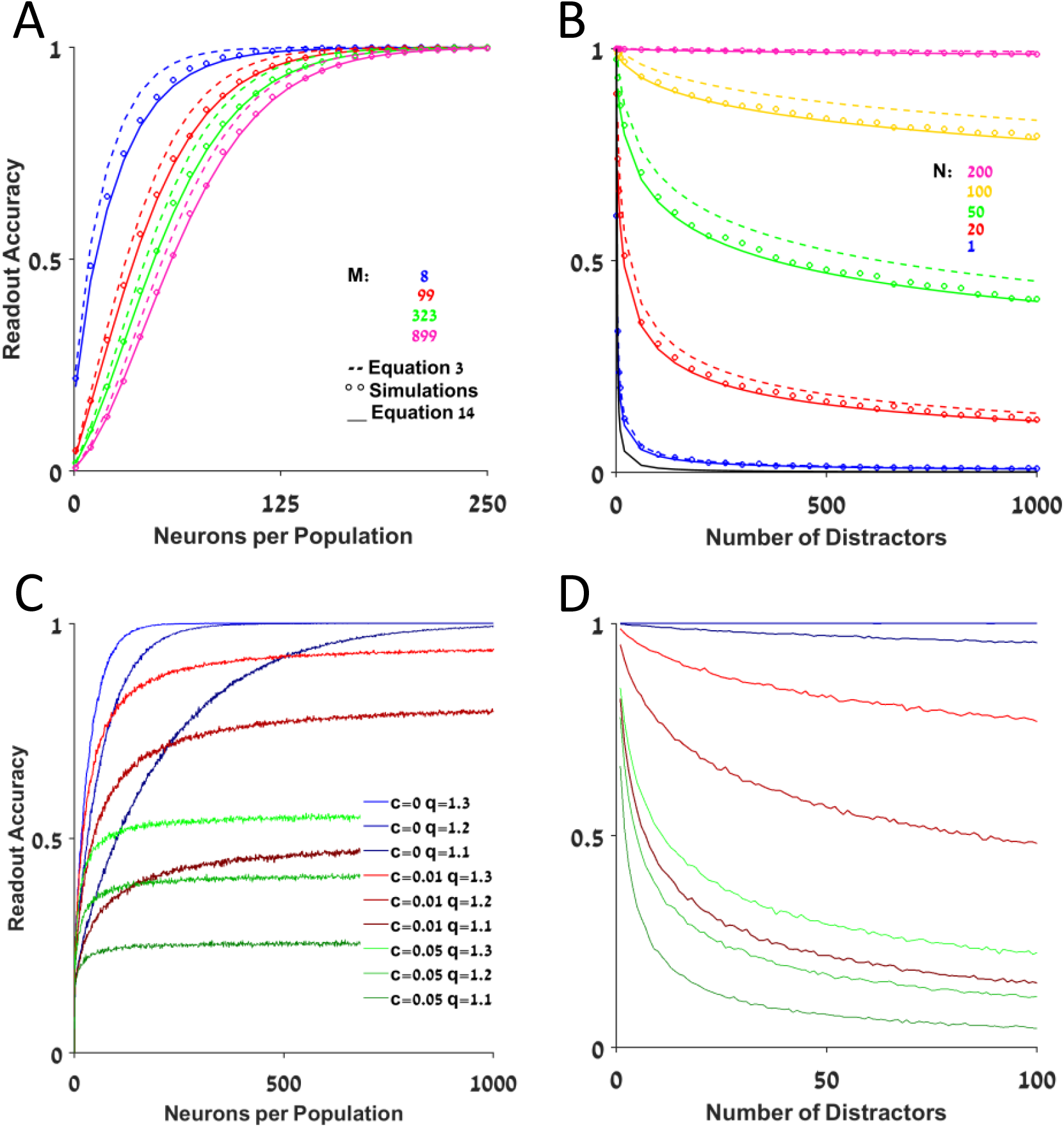
Accuracy of the Generalized WTA. (**A-D**) The accuracy of the generalized WTA is presented as a function of *N* in (A&C) and the number of distractors, *M*, in (B&D) with and without noise correlations in C-D and A-B, respectively. The open circles, dashed lines and solid lines in A-B depict the accuracy as estimated by: numerical simulations, approximation of equation 3, and approximation of equation 14, respectively. The different colors in A depict different number of distractors, *M*. The different colors in B depict different values of *N*. The colors in C-D represent different correlation levels and contextual modulation strengths.

The accuracy of the generalized WTA decays to chance value when the number of distractors is increased. In the limit of numerous distractors, *M* → ∞, we approximate (see Methods):

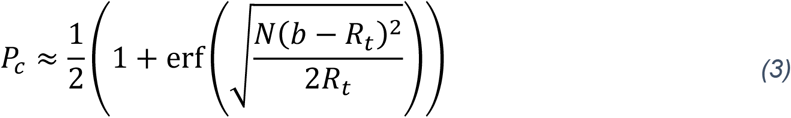

where, *R*_*t*_ is the mean firing rate of the target population, and *b* is defined as:

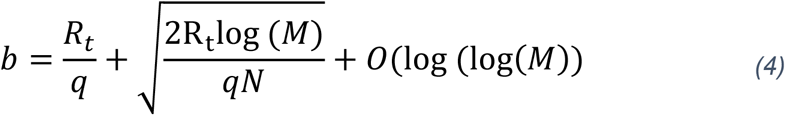

Consequently, the critical population size, *N*_c_, that reaches a certain level of accuracy *P*_*th*_ depends logarithmically on the number of distractors. Thus, to observe a deterioration in the performance of the generalized WTA, the number of distractors needs to scale exponentially in *N*. We get (see Methods):

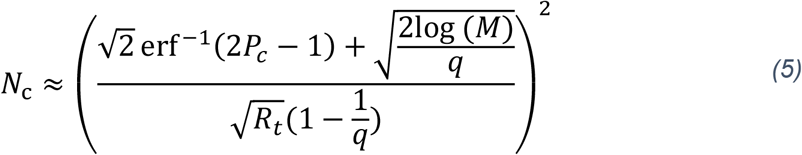

A generalized WTA readout using several hundred neurons per population can achieve an accuracy of 98% for *M* ≤ 35 (Figure 4*B*).

Heterogeneity also affects generalized WTA performance. However, averaging the neuronal responses across the population also reduces the effect of heterogeneity by a factor of 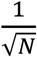. For example, compare the blue lines that correspond to different modulation strengths in Figure 4*C* to Figure 2H.

### The source of the high accuracy of the generalized WTA architecture

This remarkable improvement in the performance of the generalized WTA results from the fact that the magnitude of trial-to-trial fluctuations in the population responses was reduced by a factor of 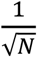due to spatial averaging. This hints that fluctuations that can generate a discrimination error are highly unlikely. However, this reasoning relies on the assumption that fluctuations in the responses of different neurons are uncorrelated. This fails to jibe with the literature reporting that noise correlations are widespread in the central nervous system [29-34] and have been a topic for extensive theoretical analyses [35-37]. It is worth noting that in contrast some studies have reported the existence of only very weak correlations [38]. Other studies have found that noise correlations tend to be stronger between pairs of neurons of similar selectivity and that there are especially strong temporal correlations in V1 [39,40], as expected in the architecture of columns we considered for the generalized case (Figure 1*E*) [41]. Thus, the effect of noise correlations on the accuracy of the generalized WTA must be considered.

### Noise correlations limit the accuracy of the generalized WTA

Our analysis showed that the accuracy of the generalized WTA in pop out tasks was undermined by noise correlations. In the limit of large populations, the generalized WTA accuracy saturates to an asymptotic value that is equivalent to the accuracy in an uncorrelated network with an effective population size of 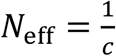, where *c* is the correlation coefficient in the neuronal responses within the same population (see Methods). Thus, noise correlations limit the ability of the generalized WTA to accumulate information from large populations of neurons. Even correlations as weak as *c* = 0.01 will cause the accuracy of the generalized WTA to saturate to *P*_c_ ≈ 90%, Figure 4*C* and 4*D*.

## Discussion

We examined the feasibility of a WTA-based algorithm as a possible neural basis for the pop out visual search task. We considered two network architectures. The first, a single best cell WTA, was based on WTA competition between contextually modulated cells. The second architecture, generalized WTA, was based on competition between populations of contextually modulated cells.

Here we focused on the scaling of the accuracy of these WTA mechanisms with population size. Our rationale was that the actual population size used in the decision is finite and limited. The population size in *V*1 can vary across different species by several orders of magnitude. Studies have shown for example that the estimated density of neurons per cubic millimeter (*mm*^3^) is approximately 160,000 in macaques [42], 40,000 in cats [43], and 180,000 in humans [44]. This makes the dependency of accuracy on the population size critically important since ceiling effects for accuracy can limit the feasibility of potential WTA mechanisms as possible explanations for pop out visual search.

We first analyzed the single best cell WTA by investigating the ability of this algorithm to correctly detect a single deviant object from the background of identical distractors. We found that the standard single-best-cell WTA accumulates information slowly as the neuronal populations size increases. The rate of improvement as a function of population size depends on the specific choice of the neuronal response distribution. WTA is sensitive to the tail of the response distribution. Thus, the accuracy in a Gaussian population is higher than in a Poisson population, which is better than an exponential population (Figure 2*A* and 2*B*).

Furthermore, when we considered the inherent neuronal heterogeneity, this slow rate of integrating information was even slower. This is because the WTA decision is determined by the extreme values of the population response, which are dominated by a small percentage of cells from out of the entire population. In addition, empirical data have indicated that the most active neurons that govern the single-best-cell WTA decision are not the most informative ones. This makes the single best cell WTA an inappropriate model when attempting to account for the observed success rate in behavioral studies.

To find a remedy for the failure of the single best cell WTA algorithm, we analyzed the generalized WTA algorithm. Here, the winning population is determined by a WTA competition between the average firing rates of each population. The analysis indicated that the generalized WTA architecture success rate was greatly improved when there was a competition between the mean (over the neural population in each column) responses. In addition, this architecture emerged as less sensitive to neuronal heterogeneity due to the spatial averaging.

However, we also found that the accuracy of the generalized WTA was limited by intra-population noise correlations but not by inter-population noise correlations. In the inter-population case, the correlations generate a collective mode of fluctuations in which the responses of all the neurons fluctuated together. Consequently, these collective fluctuations did not change the identity of the winner so that inter-population correlations had a limited effect on the accuracy.

By contrast, in the intra-population case, the correlations generated collective modes of fluctuations within the responses of the same population. Hence, these correlations affected the mean population response of separate populations differently and impacted the identity of the winner. As a result, the accuracy of the generalized WTA saturated to a size-independent limit, a highly constraining factor.

### Possible generalizations for WTA algorithms

Here we only considered a generalized WTA that utilized the mean neural response in each population. In this case, the response of each neuron had the same weight in the decision. However, due to neuronal heterogeneity, some neurons are more informative than others, such that the mean response could usefully be replaced with a weighted average. It was shown that a readout algorithm that takes this diversity into account can overcome the limiting effect of noise correlations [35,36,45]. However, this approach requires some degree of fine-tuning of the weights. Investigation of this type of algorithm is beyond the scope of the current work and will be addressed elsewhere.

Two central features that were not analyzed in the current study are the temporal aspect of the decision and the dynamical system that implements the computation. The accuracy of a temporal generalization of the WTA has been studied in the past in the framework of a two-alternative forced choice task [46]. It was shown that this generalization can yield high accuracy when required to discriminate a small number of alternatives but fails when the number of alternatives is large. This generalization can be considered as a race-to-threshold decision mechanism [47] and can be implemented by using a simple reciprocal inhibition architecture [48]. The issue of the accuracy of a dynamical implementation of a race-to-threshold WTA decision mechanism in a popout task is beyond the scope of the current work and will be addressed elsewhere.

### Temporal aspects of the decision mechanism

One parameter that cannot be well-estimated is the processing or computation time, i.e., the duration of information integration from the contextually modulated cells. Longer processing times make the single neuron response more informative, since both the mean and the variance of the spike count are expected to scale linearly with the processing time. As a result, the accuracy of both WTA architectures will also improve.

What is a reasonable estimate for processing time? One can bound processing time from above by the reaction time, which is on the order of 500 milliseconds and includes both the delay of the sensory response and the motor output. A tighter estimate of processing time was obtained by Stanford and Salinas in a two-alternative forced choice task [49]. Using a cleverly designed experiment, they estimated processing time to be on the order of a few tens of milliseconds. Here, we used a conservative estimate of 200 ms for the processing time. Shorter durations will only further decrease the accuracy of the WTA.

## Conclusion

Overall, the two architectures are limited in their ability to account for the high accuracy observed in behavioral studies. This analysis highlights the fact that the high accuracy observed experimentally in pop out tasks is not straightforward and requires further theoretical and experimental considerations.

## Methods

### WTA accuracy

In a homogeneous network, assuming the responses of different neurons are independent, given the stimulus, the accuracy of the WTA is given by:

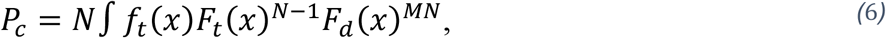

where *f*_*α*_(*x*), α= *t, d* is the probability density of the neural response *x* in a target (*α* = *t*) or distractor (*α* = *d*) populations, and *F*_*α*_(*x*) is the cumulative distribution function.

### Exponential response distribution

The case of an exponential response distribution, 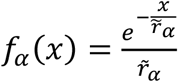, is more amenable to analytical treatment. In this case Eq (6) yields:

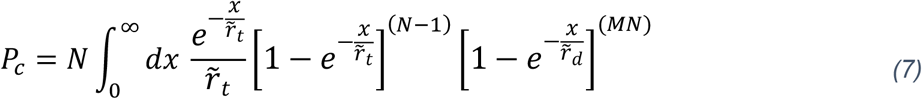

#### The case of N=1

For the simple case of one neuron per population, *N* = 1, the WTA accuracy is:

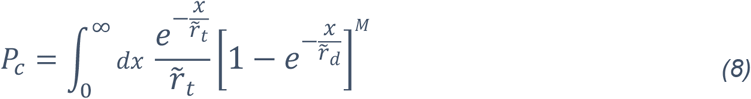

By the change of variable 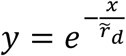, one obtains:

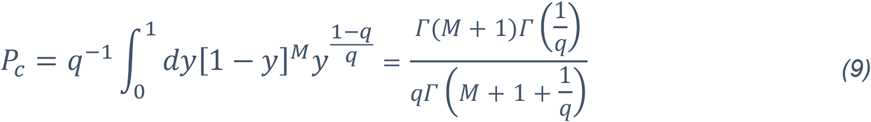

In the limit of *q* → 1, *P*_c_ converges to chance value, 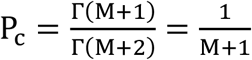. To study the limit of large *q* for finite *M* we used the first order expansion of the gamma function: 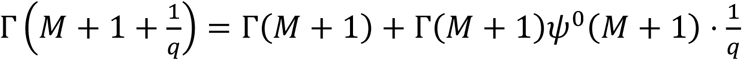,where ψ^0^ is the zero order of the polygamma function. Next, we expanded the polygamma function as 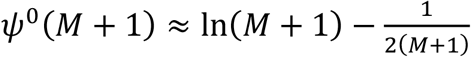 . Using the above approximation yields 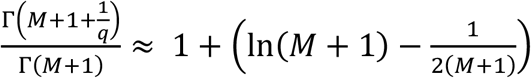. For large *q* we can approximate 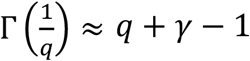, where *γ* is the Euler-Mascheroni constant, obtaining:

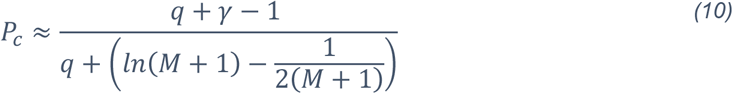

For finite *M, P*_*c*_ approaches 1 algebraically in *q*.

For *M* ≫ 1, by using the asymptotic expansion of the gamma function one obtains 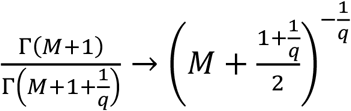, which yields:

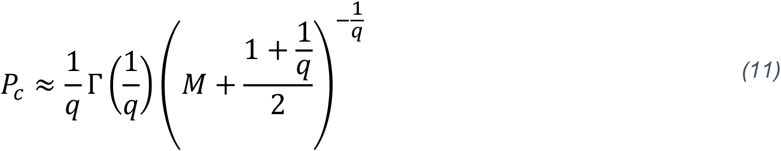

Thus, for any fixed *q* > 1, *P*_*c*_ decays to chance algebraically in *M*.

**The case of N>1**

Changing variables to y ≡ 1− *F*_*t*_(*x*) in equation 6 yields:

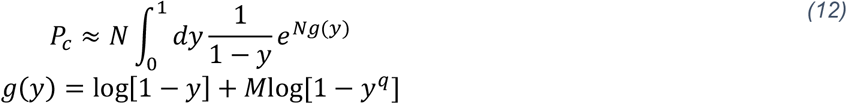

For large *N*, this integral is dominated by small y. Substituting *u* =−*g*(y), and noting that for small y, (*i*) *u* ≈y + *M*y^*q*^ (for finite *M* in the limit of large *N*), and (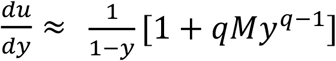we obtain

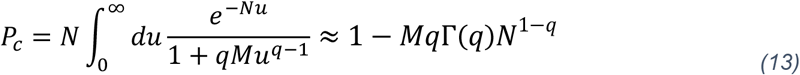

where we used Watson’s lemma to obtain the last result.

### Participation rate of a neuron

The participation rate of neuron *i* in population *j* is the probability of neuron *i* to be the winner given that the *j*′*th* population won.

### Generalized WTA accuracy

In this generalized WTA competition, the stimulus is estimated by the receptive field of the most active population. The activity of population *α∈{*1, … *M* + 1} is given by 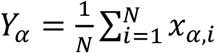, where *x*_α*i*,_ is the response (number of spikes) of neuron *i* of population *α*. Assuming the responses of different neurons are statistically independent, given the stimulus, in the limit of large *N*, we can approximate the population activities by independent Gaussian random variables with 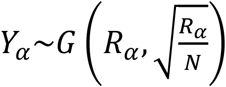, where 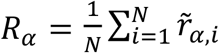where 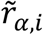is the trial to trial average of the response of neuron *i* of population *α*. In addition, we selected that the variance of the single neuron response is equal to its mean.

In realistic networks, the trial-to-trial average activity of population *α, R*_*α*_, fluctuates from one population to another due to the inherent neuronal heterogeneity with mean ≪ *R*_*α*_ ≫=*r*_*x*_ (*x* =t, *d* for target and distractor populations, respectively) and variance 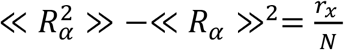, where we define ≪ ≫, as averaging with respect to the quenched disorder i.e the neuronal heterogeneity. For large *N*, the fluctuations in *R*_*α*_ become negligible relative to its mean and we can ignore the fluctuations in *R*_*α*_.

For a sufficiently large number of distractors, *M*, we can use the central limit for extreme values for Gaussian distribution to approximate the cumulative distribution of the maximal value of activity of the *M* distractor populations by a Gumble distribution, yielding:

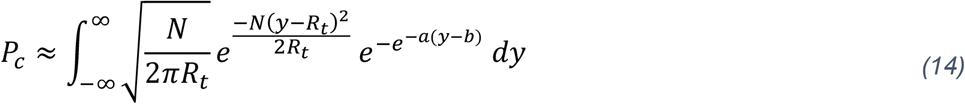

With 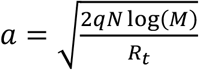, and 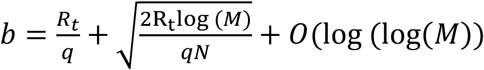.

### Approximation of Generalized WTA accuracy

The argument in the integral of equation 14 consists of a Gaussian probability density multiplied by a Gumble distribution. For large *M*, we approximate the Gumble distribution by a Heaviside function centered around *b*, yielding:

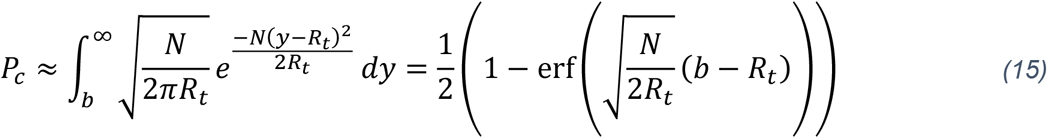

Thus, for any fixed number of distractors the accuracy of the generalized WTA will converge to 1 exponentially in *N*.

### Derivation of *Nceiling*

The typical number of neurons required to obtain a certain degree of accuracy, *P*_*th*_, denoted *N*_*c*_ can be derived from equation 16, yielding

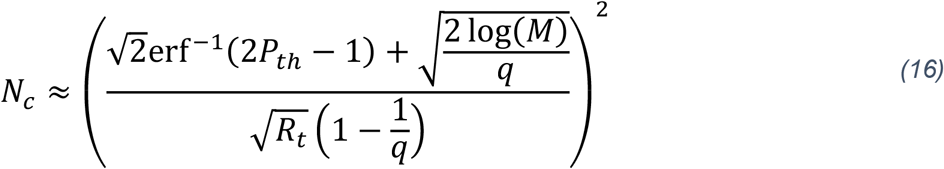

As expected, for *q* → 1 or for *P*_*th*_ → 1, one obtains *N*_*c*_ → ∞. From Equation (16) one finds that *N*_*c*_ scales logarithmically with the number of distractors, *M*.

### Effect of correlations on the Generalized WTA

To study the effect of noise correlations on the accuracy of the generalized WTA, we modeled the response statistics of the network by multivariate Gaussian distribution. Thus, the neural responses *{x*_*α,i*_} (*α* ∈ *{*1, … *M* + 1}, *i* ∈ *{*1, … *N*}) are Gaussian random variables with means 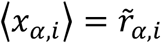and covariance 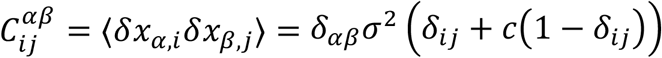, where *σ*^2^ is the variance of the trial-to-trial fluctuations in the single neuron response, and *c* is the pairwise correlation coefficient of the responses of neurons within the same population. Correlations redistribute the noise across various dimensions of the neural response space [50]. Here we focused on uniform correlations within populations since these specific correlations are particularly pertinent to constraining the accuracy of the generalized WTA.

For large *N*, neglecting the fluctuations due to neuronal heterogeneity the population activities are independent Gaussian random variables with means 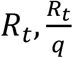and variances 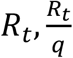, for the target, and distractor populations respectively. Thus, the accuracy of network with within-population correlation coefficient *c* is equal to that of an uncorrelated network of size 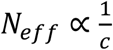.

### WTA error for different noise types

Decision errors of the WTA algorithm result from large fluctuations in the neural responses. The likelihood of these fluctuations depends on the tail of the response distribution. The tail of the distribution behaves differently for exponential, *P*(*x*) *∼ e*^−*xr*^, Poisson, *P*(*x*) *∼ e*^−*x*·*ln*(*x*)^, and Gaussian, 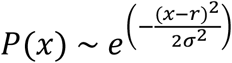Consequently, for a similar mean and variance, the WTA algorithm is expected to perform better for an exponential population of neurons than for Poisson, and for Poisson better than for Gaussian. Compare for example the red and blue traces in Figures 2*A*−*B*.

## Numerical methods

### Realistic network architecture

The dataset reported in Ben-Tov and colleagues [17] contained 65 neurons, out of which we selected 23 contextually modulated neurons. Neurons whose estimated mean firing rate in the pop out condition minus two times the standard error of its mean were greater than the estimated mean firing rate in the uniform condition plus two times the standard error of this mean were selected.

Each neuron in the dataset was characterized by two parameters: its mean firing rate in response to a pop out stimulus and its contextual modulation strength. The mean firing rate of the neuronal population to pop out and uniform stimuli was 12.9 H*z* and 8.9 H*z*, respectively, and the mean contextual modulation strength *q* was 1.44 (see figure 2*D*).

However, the firing rates of individual neurons were widely distributed around the population average, Figure 2*D*.

To generate a realization of an individual network consisting of (*M* + 1) populations of *N* neurons each, we drew *N* × (*M* + 1) neurons from the set of 23 contextually modulated cells randomly with equal probabilities and with replacement. To generate a single trial of the neural response to a pop out stimulus, the deviant population was chosen first. Then, the spike count for each neuron was drawn independently from either a Poisson or an exponential distribution. A time window of 200 [*ms*] [49] was used to translate the mean firing rate of each neuron to the mean spike count response to the stimulus.

### Artificial networks

To generate a network of *M* + 1 populations with *N* neurons each, we drew the firing rates in response to target stimulus, 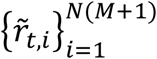, and contextual modulation strengths, 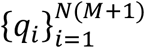, for each of the *N* · (*M* + 1) neurons randomly in an independent manner. Specifically, the firing rates in response to a pop out stimulus were drawn from a log-normal distribution [51,52] with mean *μ* = 12.8, and standard deviation *σ* = 3.57. The contextual modulation strengths were drawn from an exponential distribution, with mean 1.44, as in the empirical data. The accuracy of the WTA in a single realization of an individual (represented by a single network of *N* · (*M* + 1) neurons) was estimated by simulating the network response in 1000 trials.

### Parameters for the numerical simulations

| Figure 2 panel | $N$ | $M$ | $r_t$ | $q$ | $Trials$ | $Realizations$ |
| --- | --- | --- | --- | --- | --- | --- |
| A | - | 48 | 12.8 | 1.44 | 1000 | 20 |
| B | 100 | - | 12.8 | 1.44 | 1000 | 20 |
| C | - | - | 12.8 | 1.44 | 1000 | 20 |
| D | - | 48 | $\log(x) \sim N(12.8, 12.8)$ | $x \sim \exp(\lambda = 1.44)$ | 1000 | 100 |
| E | 100 | - | $\log(x) \sim N(12.8, 12.8)$ | $x \sim \exp(\lambda = 1.44)$ | 1000 | 100 |
| G | 100 | 48 | $\log(x) \sim N(12.8, 12.8)$ | $x \sim \exp(\lambda = 1.44)$ | 10000 | 1 |
| H | - | 8 | $\log(x) \sim N(12.8, 12.8)$ | $x \sim \exp(\lambda = 1.44)$ | 1000 | 20 |
| I | 5000 | - | $\log(x) \sim N(12.8, 12.8)$ | $x \sim \exp(\lambda = 1.44)$ | 1000 | 20 |
| Figure 3 panel |  |  |  |  |  |  |
| A | 100 | 8 | $\log(x) \sim N(12.8, 12.8)$ | $x \sim \exp(\lambda = 1.44)$ | 1000 | 1 |
| B | 100 | 8 | $\log(x) \sim N(12.8, 12.8)$ | $x \sim \exp(\lambda = 1.44)$ | 1000 | 1 |
| C | 100 | 8 | $\log(x) \sim N(12.8, 12.8)$ | $x \sim \exp(\lambda = 1.44)$ | 1000 | 20 |
| D | 100 | 8 | $\log(x) \sim N(12.8, 12.8)$ | $x \sim \exp(\lambda = 1.44)$ | 1000 | 1 |
| E | 100 | 8 | $\log(x) \sim N(12.8, 12.8)$ | $x \sim \exp(\lambda = 1.44)$ | 1000 | 1 |
| F | 1000 | 8 | $\log(x) \sim N(12.8, 12.8)$ | $x \sim \exp(\lambda = 1.44)$ | 1000 | 1 |
| G | 1000 | 8 | $\log(x) \sim N(12.8, 12.8)$ | $x \sim \exp(\lambda = 1.44)$ | 1000 | 1 |
| H | - | 8 | $\log(x) \sim N(12.8, 12.8)$ | $x \sim \exp(\lambda = 1.44)$ | 1000 | 20 |
| I | 1000 | 8 | $\log(x) \sim N(12.8, 12.8)$ | $x \sim \exp(\lambda = 1.44)$ | 1000 | 1 |
| J | 1000 | 8 | $\log(x) \sim N(12.8, 12.8)$ | $x \sim \exp(\lambda = 1.44)$ | 1000 | 1 |
| Figure 4 panel |  |  |  |  |  |  |
| A | - | - | $\log(x) \sim N(12.8, 12.8)$ | $x \sim \exp(\lambda = 1.44)$ | 1000 | 50 |
| B | - | - | $\log(x) \sim N(12.8, 12.8)$ | $x \sim \exp(\lambda = 1.44)$ | 1000 | 50 |
| C | - | 8 | $\log(x) \sim N(12.8, 12.8)$ | $x \sim \exp(\lambda = 1.44)$ | 1000 | 50 |
| D | 100 | - | $\log(x) \sim N(12.8, 12.8)$ | $x \sim \exp(\lambda = 1.44)$ | 1000 | 50 |

## Acknowledgments

This work has been supported in part by the Israel Science Foundation grants no. 824/21 and 624/22.

